# A novel quantification method for RPE phagocytosis using a VLC-PUFA-based strategy

**DOI:** 10.1101/2022.07.02.498584

**Authors:** Fangyuan Gao, Emily Tom, Stephanie A. Lieffrig, Silvia C. Finnemann, Dorota Skowronska-Krawczyk

## Abstract

The vertebrate retinal pigment epithelium (RPE) lies adjacent to the photoreceptors and is responsible for the engulfment and degradation of shed photoreceptor outer segment fragments (POS) through receptor-mediated phagocytosis. Phagocytosis of POS is critical for maintaining photoreceptor function and is a key indicator of RPE functionality. Popular established methods to assess RPE phagocytosis rely mainly on quantifying POS proteins, especially their most abundant protein rhodopsin, or on fluorescent dye conjugation of bulk, unspecified POS components. While these approaches are practical and quantitative, they fail to assess the fate of POS lipids, which make up about 50% of POS by weight and whose processing is essential for life-long functionality of RPE and retina. Here, we have developed a novel VLC-PUFA-based approach for evaluating RPE phagocytic activity by primary bovine and rat RPE and the human ARPE-19 cell line and validated its results using traditional methods. This new approach can be used to detect the dynamic process of phagocytosis at varying POS concentrations and incubation times and offers a robust, unbiased, and reproducible assay that will have utility in studies of POS lipid processing.

## INTRODUCTION

Phagocytosis of photoreceptor outer segment (POS) by the retinal pigment epithelium (RPE) is essential for retinal health and function. Disruptions in phagocytosis have been previously linked to retinal degeneration in inherited retinal diseases, aging, and age-related macular degeneration (AMD)(1-3). Due to the importance of this process in maintaining visual function, it has remained an area of intense study, and several techniques, both *in situ* and *in vitro*, have been developed over the past 50 years to quantify RPE phagocytosis. Common *in situ* methods to study RPE phagocytosis have been recently reviewed (4). Briefly, light microscopy(5), transmission electron microscopy (TEM)(6), and immuno-electron microscopy (7) allow for visualization of phagosomes within RPE. While *in situ* methods provide valuable insight into candidate proteins that may be important in POS renewal in animal models such as mice with different mutations, evaluating POS uptake by cultured RPE cells *in vitro* possesses several potential applications and advantages: 1) Defects in RPE or photoreceptor function during the normally synchronized process of phagocytosis can be separated from one another. 2) Depending on the gene or protein of interest, different phases of the phagocytic process, namely particle recognition/binding, internalization, and digestion, can be distinguished (8). 3) Using genetic or pharmacological approaches, RPE cells can be manipulated to simulate an *in vivo* RPE cell-specific disease state(9). 4) High-throughput screening platforms can be established for screening chemical and biological libraries to identify potential compounds that can rescue phagocytosis deficiencies in RPE (10). The current methods to quantify POS uptake by RPE cells in culture include fluorescence-based assays, such as fluorescence-activated cell sorting (FACS) (11), fluorescence scanning (12), or immunofluorescence (8), which require POS to be first labeled with a fluorescent dye such as FITC, and immunoblot-based assays, where unlabeled POS can be detected using an antibody specific to POS, such as transducin (13) or rhodopsin (9).

As different components of the POS are processed at different rates in POS phagolysosomes, single antigen immunodetection is not representative for the entirety of POS components. For example, different monoclonal antibodies to rhodopsin recognize either the C- or N-terminus of rhodopsin, and these epitopes are lost from immunolabeling at different rates due to their varying stability in RPE phagolysosomal digestion (14). Moreover, loading RPE cells in culture with the lipofuscin component A2E has no effect on rhodopsin processing, but slows degradation of FITC-POS. This implies that lysosomal degradation of non-rhodopsin POS components is affected; however, the precise nature of such components remain unknown (15). Currently, established methods do not assess levels and rates of processing of POS lipids, which comprise about 50% of POS. Altogether, there is thus a critical need for additional methodology for POS quantification that is quantitative, and lipid focused to complement existing assays.

In the retina, very-long-chain polyunsaturated fatty acids (VLC-PUFAs, which contain acyl chains longer than 26 carbons) are produced in photoreceptors through the elongation of LC-PUFAs mediated by Elongation of very-long-chain fatty acids like-4 (ELOVL4) and are highly enriched in the light-sensitive membrane disks of the photoreceptor outer segments (15, 16). Following shedding and subsequent phagocytosis of POS by RPE cells, photoreceptor-derived VLC-PUFAs are efficiently recycled back to the inner segments of photoreceptors for further use (17).

Here, we present direct evidence that RPE cells do not produce or maintain VLC-PUFAs endogenously. Based on this finding, we developed a novel phagocytosis assay based on the quantification of phagocytosed VLC-PUFAs by RPE cells. The proposed VLC-PUFA-based assay has the ability to quantify phagocytosis in RPE cells in culture after incubation with different concentrations of POS and different incubation times. This new method is also suitable for the quantification of defects in RPE phagocytic activity. Results generated using this VLC-PUFA-based assay are complementary to traditional immunofluorescence- and immunoblot-based methods, providing a lipid-based approach for rapid evaluation of POS phagocytosis by RPE.

## MATERIALS AND METHODS

### Animals

Pink-eyed dystrophic Royal College of Surgeons (RCS) rats (rdy/rdy-p) originally obtained from the National Center for Research Resources (NIH, Bethesda, MD) and Sprague-Dawley (SD) wild-type (WT) albino rats originally obtained from Charles River Labs (Wilmington, MA) were raised in a 12-h light/12-h dark light cycle with standard food and water ad libitum and bred to yield litters for RPE isolation. Animal experimentation was conducted according to the ARVO guidelines for the Use of Animals in Ophthalmic and Vision Research and reviewed and approved by the Institutional Animal Care and Use Committee of Fordham University.

### Cell culture and siRNA transfection

ARPE-19 cells were obtained from ATCC and maintained in DMEM/F-12 containing 10% fetal bovine serum (FBS). Primary bovine retinal pigment epithelial cells (RPE) were generated by Dr. Huajun Yan and maintained in RtEBM (Lonza, USA) supplemented with RtEGM SingleQuots Supplement Pack (Lonza, USA). Both cell lines were incubated at 37 °C with 5% CO_2_. ARPE-19 cells and primary bovine RPE cells were seeded in 12-well plates and grown to full confluency before experiments.

ARPE-19 cells were electroporated with three unique siRNA specific for human *MERTK*, *GAS6*, and *MFGE8* (Table 1, Integrated DNA Technologies (IDT), USA), and cells electroporated without siRNA was used as a negative control, using Neon™ Transfection System (Thermo Fisher, USA) according to the manufacturer’s protocol. Experiments were carried out 5 days after the transfection procedure. The effectiveness of the *MERTK*, *GAS6*, and *MFGE8* knockdown was examined using quantitative RT-PCR (qRT-PCR).

**Table 1.**
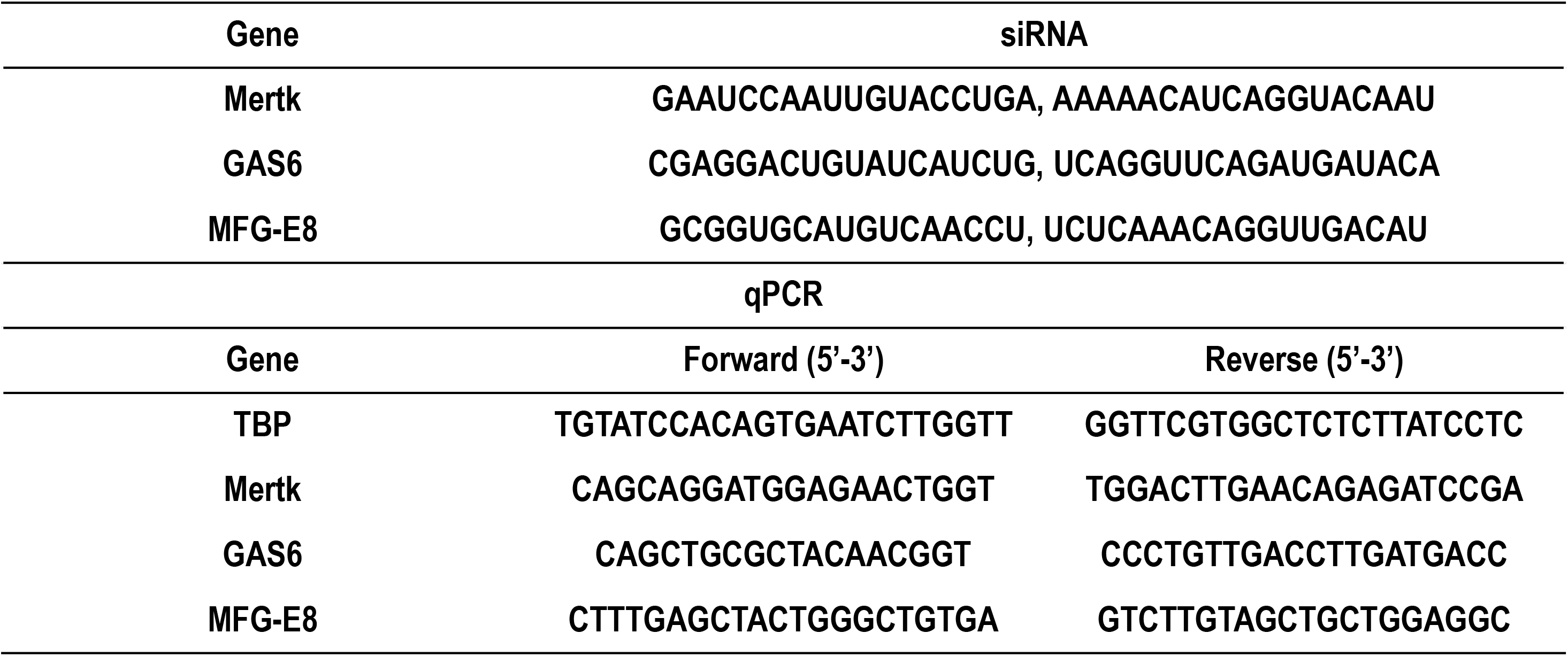
siRNA and qPCR primer sequences.

Rat primary RPE cells were isolated from 4–9-day-old SD and RCS rats as described previously (18). Mixed sex litters were used. Briefly, the lens and vitreous body were removed from freshly enucleated eyes, and eyecups were dissected following sequential hyaluronidase and trypsin enzymatic treatments. Sheets of RPE were manually collected and re-trypsinized for 90s. Cells were cultured in DMEM with 10% FBS for 4 days before experiments.

### RNA Extraction, Reverse Transcription, and qPCR Analysis

Total RNA was extracted from cells using RNeasy Plus mini kits (Qiagen). RNA concentration was measured by a Qubit fluorescence assay (Thermo Fisher, USA). A total 1 μg of extracted RNA was reverse transcribed using SuperScript IV VILO Master Mix kit (Thermo Fisher, USA), according to manufacturer’s instructions. cDNA was diluted 1:10 in nuclease free water prior to qPCR analysis using a CFX384 real-time PCR system (Bio-Rad, USA). Reaction mixtures containing 5 μl PowerUp SYBR Green Master Mix (Applied Biosystems, USA), 4 μl cDNA, 1 μl primer mix (final concentration of 200 nM per primer) were prepared in Hard-Shell PCR 384-well plates (Bio-Rad, USA). Amplification conditions were as follows: 50 °C for 2 min, followed by 40 cycles at 95 °C for 15 s and 60 °C for 60 s. Melting curve analysis was performed from 65–95 °C. All samples were run in triplicate and mean Ct (cycle threshold) values were used for further analysis.

### Lipid purification and fatty acid extraction from RPE cells and tissue

Cell pellets were resuspended in 200 μL ddH_2_O and transferred into a clean glass test tube. 1 μg 6Z,9Z,12Z,15Z,18Z-heneicosapentaenoic acid (FA 21:5, Cayman, USA) was added as internal standard. Lipids were extracted by the Bligh-Dyer method followed by hydrolysis and extraction of total fatty acids. Briefly, 750 μL ice-cold 1:2 (v/v) CHCl_3_:MeOH was added directly to the sample and the sample was vortexed. Then, 250 μL CHCl_3_ was added and mixed well. 250 μL ddH_2_O was added and vortexed well to produce a two-phase system. After centrifuging at 3000 RPM for 3 min, the bottom phase was collected and evaporated under a constant stream of nitrogen. To recover the total fatty acids, 720 μL acetonitrile and 10 μL hydrochloric acid were added to the dried film and the sample was kept at 95 °C for 1 h. Then, 1 ml of hexane was added to the sample and vortexed. The upper phase containing total fatty acids was transferred to a new tube and evaporated under nitrogen. The total fatty acids were dissolved in acetonitrile-isopropanol (50:50, v/v) for further LC-MS Analysis (**Fig. 1A**).

**Figure 1.**
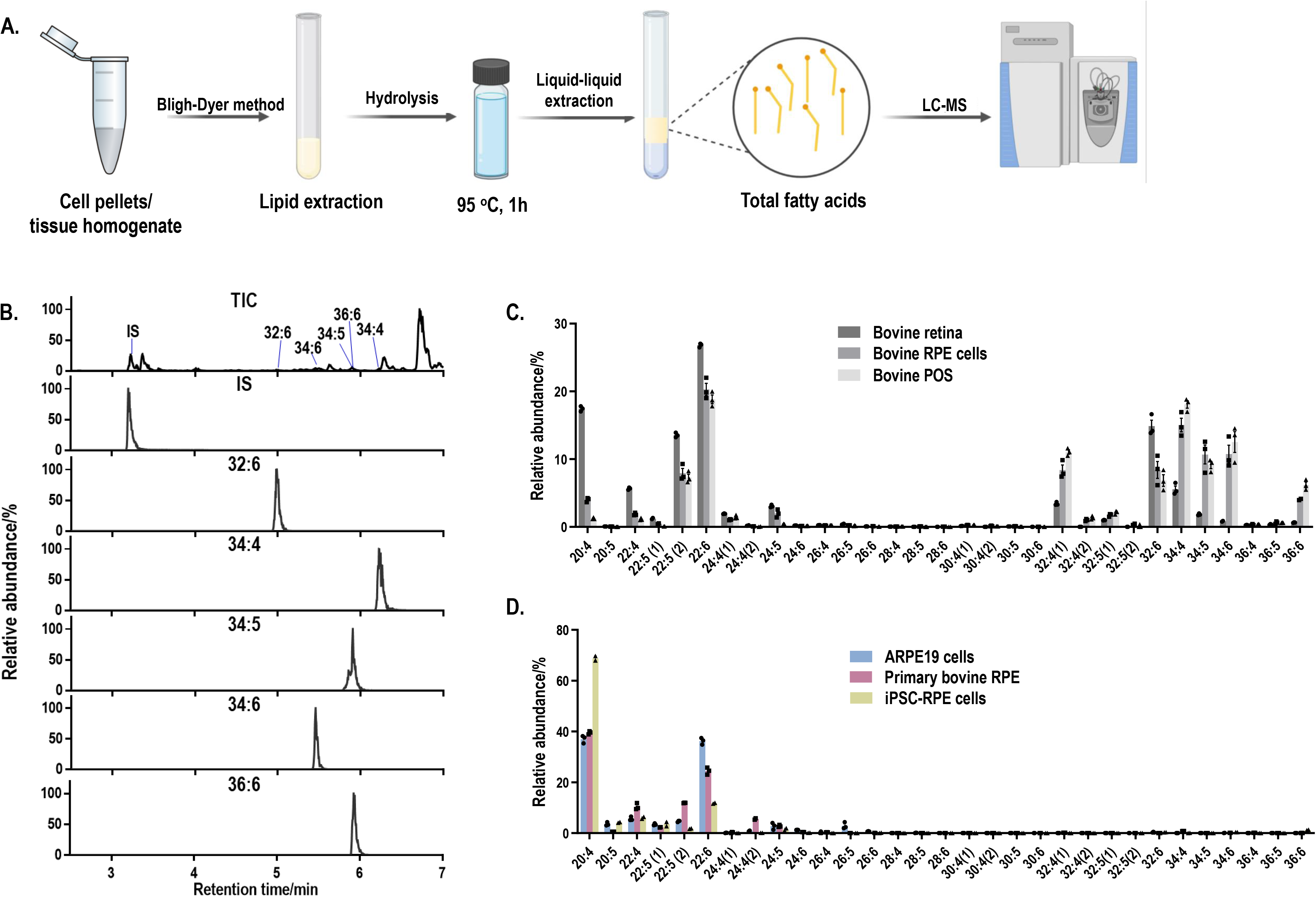
LC-MS method to detect LC-PUFAs and VLC-PUFAs. **A.** Workflow for lipid purification and fatty acid extraction from cells and tissue. **B.** Total ion (TIC) and extracted ion (EIC) mass chromatograms of VLC-PUFAs obtained in full scan (FS) mode by LC–ESI(-)–MS (FS). IS: internal standard (21:5) **C.** Abundance plots present the relative abundances within the total abundance of all the detected LC-PUFAs and VLC-PUFAs in bovine samples. LC-PUFAs and VLC-PUFAs composition in bovine RPE and retina tissues exhibited high abundance of VLC-PUFAs, ranging from 32:4 to 36:6. **D.** Abundance plots present the relative abundances within the total abundance of all the detected LC-PUFAs and VLC-PUFAs in RPE cells. LC-PUFAs and VLC-PUFAs composition in cultured RPE cells showed minimal amounts of endogenous VLC-PUFAs

Tissue samples were homogenized in ddH_2_O. 1 μg FA 21:5 was added as internal standard. Lipids were extracted by the Bligh-Dyer method, followed by hydrolysis and extraction of total fatty acids as described above.

### LC-MS Analysis of VLC-PUFAs

Separation of VLC-PUFAs was achieved on an Acquity UPLC® BEH C18 column (1.7 μm, 2.1 mm × 100 mm, Waters Corporation) using a mobile phase consisting of 1% 1 mol/L ammonium acetate and 0.1% formic acid in water (A) acetonitrile/isopropanol (1:1, v/v), 1% 1 mol/L ammonium acetate, and 0.1% formic acid (B) at a flow rate of 0.4 mL/min, with the following linear gradient: 0-2 min, 35% -80% B, 2-7 min, 80-100% B,7-14 min, 100% B, 14-15,100-35% B.

For VLC-PUFAs analyses, 14Z,17Z,20Z,23Z,26Z,29Z-dotriacontahexaenoic acid (32:6, Cayman, USA) and the extracted mixed VLC-PUFAs from bovine retina was used as VLC-PUFA standards, since most commercial VLC-PUFA standards are not available, and retina contains the highest levels of VLC-PUFAs of any mammalian tissue analyzed thus far (20-22). The identification of each VLC-PUFA in retina samples was conducted extracting a chromatographic peak for each of the corresponding [M-H]^-^ ions using 5 ppm tolerance in full scan mode. Then fragments of the different VLC-PUFAs were studied through a separated parallel reaction monitoring (PRM) method for further identification of the compounds.

The established retention times were used for identification of LC-PUFAs and VLC-PUFAs in phagocytosed POS (**Table 2**). For quantification, the Q Exactive MS (Thermo Fisher Scientific, Waltham, MA) was operated in a full MS scan mode (resolution 70,000) in negative mode with a scan range of m/z 250–800, and the VLC-PUFAs of interest were quantified by extracting a chromatographic peak for the corresponding [M-H]^-^ ions using 5 ppm tolerance. The comparison was made by normalizing each peak area with the internal standard fatty acid 21:5.

**Table 2.** m/z and retention time of LC-PUFA and VLC-PUFA from standard solution and bovine retina.

| FAs | m/z | Retention time/min |  |
| --- | --- | --- | --- |
|  |  | Standard solution | Bovine retina |
| 21:5(IS) | 315.233 |  | 3.19 |
| 20:4 | 303.233 |  | 3.3 |
| 20:5 | 301.2173 |  | 3.11 |
| 22:4 | 331.2643 |  | 3.58 |
| 22:5 (1) | 329.2486 |  | 3.35 |
| 22:5 (2) | 329.2486 |  | 3.44 |
| 22:6 | 327.233 | 3.24 | 3.23 |
| 24:4(1) | 359.2956 |  | 3.91 |
| 24:4(2) |  |  | 4.02 |
| 24:5 | 357.2799 | 3.66 | 3.69 |
| 24:6 | 355.2643 |  | 3.45 |
| 26:4 | 387.3269 |  | 4.32 |
| 26:5 | 385.3112 |  | 4.03 |
| 26:6 | 383.2956 |  | 3.73 |
| 28:4 | 415.3582 |  | 4.76 |
| 28:5 | 413.3425 |  | 4.42 |
| 28:6 | 411.3269 |  | 4.12 |
| 30:4 | 443.3895 |  | 5.26 |
| 30:5 (1) | 441.3738 |  | 4.89 |
| 30:5 (2) | 441.3738 |  | 4.99 |
| 30:6 | 439.3582 |  | nd |
| 32:4 (1) | 471.4208 |  | 5.76 |
| 32:4 (2) | 471.4208 |  | 5.91 |
| 32:5 (1) | 469.4051 |  | 5.34 |
| 32:5 (2) | 469.4051 |  | 5.45 |
| 32:6 | 467.3895 | 4.98 | 5.01 |
| 34:4 | 499.4521 |  | 6.28 |
| 34:5 | 497.4364 | 5.88 | 5.89 |
| 34:6 | 495.4208 | 5.42 | 5.46 |
| 36:4 | 527.4834 |  | 6.67 |
| 36:5 | 525.4677 |  | 6.34 |
| 36:6 | 523.4521 |  | 5.94 |

ND = non-detected

### POS Phagocytosis Assay

RPE cells were seeded at equal density on 12-well plates to have at least triplicate samples for each condition. Then, the RPE cells were cultured in the cell culture incubator until they were fully confluent.

Bovine POS (InVision BioResources) was counted and resuspended in cell culture medium. POS aliquots were stored at -80 °C. For phagocytosis assays, cell number was determined, and appropriate POS concentration (POS/cell) was added. RPE cells were incubated with POS in the cell culture incubator for the appropriate time. Phagocytosis was terminated by washing 3 times with PBS.

### Quantification of phagocytosed POS by VLC-PUFA-based assay

To detect internalized POS only by VLC-PUFA-based assay, trypsin containing EDTA (0.25%) was added to RPE cells and incubated in the cell culture incubator until the cells were transparent, but still attached. Trypsin-EDTA, along with most of the surface-bound POS, was removed and the cells were then harvested and washed with PBS for 3 times. The cell pellet was kept at -20°C for further analysis.

For quantification of phagocytosed POS, the lipids were extracted and hydrolyzed. VLC-PUFAs 32:6, 34:4, 34:5, 34:6, 36:6 were analyzed as the markers of internalized POS following the procedures mentioned above.

### Quantification of phagocytosed POS by immunocytochemical analysis

Immunofluorescence quantification of POS phagocytosis was performed following a previously established protocol (8). Briefly, confluent ARPE-19 monolayers grown on coverslips were incubated with 5, 10, or 20 FITC-conjugated POS/cell for 2 and 5 hours. Washed cells were briefly fixed in 4% PFA in PBS for 20 min and the remaining fixative was quenched with NH_4_Cl. Cells were blocked in 1% BSA for 10 min and incubated with mouse-rhodopsin antibody (clone 1D4, a kind gift from Dr. David Salom) diluted 1:1000 in 1% BSA for 25 min. After washing 3 times in PBS, cells were incubated with goat anti-mouse AlexaFluor-647 secondary antibody diluted 1:5000 in 1% BSA for 1 hour. After washing 3 times in PBS, nuclei were stained with DAPI, and coverslips were mounted with Fluoromount G. Images were acquired using Keyence All-in-One Fluorescence Microscope (BZ-X800, Osaka, Japan) at 100X magnification, and the number of green particles was counted and normalized to the number of cell nuclei using single extract colocalization in Keyence Hybrid Cell Count software.

### Quantification of phagocytosed POS by Western Blot

Opsin immunoblotting was performed following a previously established protocol (8). Confluent ARPE-19 monolayers were incubated with 10 POS/cell for 2 hours. Triplicate samples were designated for detection of either total (bound plus internal) POS or only internalized POS. Samples designated for total POS detection were washed with PBS-CM (PBS with calcium and magnesium), while only internalized POS samples were incubated with PBS-EDTA for 10 minutes. Total protein was extracted with ice-cold lysis buffer (20mM HEPES pH 7.9, 420mM NaCl, 3mM MgCl_2_, 10% glycerol with freshly supplemented 5mM DTT and protease inhibitor cocktail), subjected to SDS-PAGE electrophoresis, transferred onto a PVDF membrane, and blocked with casein in PBS for 1 hour. The membrane was incubated with mouse-rhodopsin (clone 1D4) antibody (1:1000) overnight at 4 °C. Following 3 washes with PBST, the membrane was incubated with HRP-conjugated secondary antibodies for 1 hour. The bands were visualized by a chemiluminescent reaction and imaged using the ChemiDoc Imaging System. Densitometric analysis of obtained bands was performed by FIJI, and the data were normalized to total protein stained by Ponceau S.

### Phagocytosis by primary rat RPE cells

POS phagocytosis assays were adapted from published protocols (19). Confluent, primary rat wild-type or MERTK-deficient RCS RPE cells in wells of a 48-well plate were challenged with purified POS (∼10 POS/cell) in the presence of 1 µg/mL recombinant mouse MFG-E8 and 2 µg/mL human protein S in DMEM at 37°C for 45 min, followed by removal of excess POS, rinse with DMEM, and incubation in DMEM with 2 µg/mL protein S for 1 h to promote further POS internalization. Cells were washed three times with ice-cold PBS, trypsinized for 5 min at 37°C to remove bound POS, resuspended in PBS and pelleted at 500 x g for 5 min at 4°C. Pellets were resuspended and washed twice with PBS. After the final rinse, 10% of the cell suspension was separated for DNA analysis and 3% was separated for Western blot protein analysis. Remaining cells were pelleted, and dry cell pellets were stored at -80°C until further Western Blot or lipid analysis.

For immunoblotting, cell suspension aliquots were lysed in RIPA buffer supplemented with protease inhibitor cocktail. Proteins were separated by reducing SDS-PAGE on 4-20% gradient polyacrylamide gels, transferred to nitrocellulose membranes, and blocked in 10% non-fat milk powder in TBS. Blots were incubated with rhodopsin (clone B6-30, a kind gift from Dr. Paul Hargrave (20) and alpha-tubulin (Cell Signaling CAT #2125) primary antibodies, and incubated with horseradish peroxidase-conjugated secondary antibodies followed by enhanced chemiluminescence digital detection using a KwikQuant Imager. Bands were quantified by densitometry using ImageJ software.

## RESULTS

### VLC-PUFA quantification - method development

To obtain the PUFA composition from different samples, we developed a Liquid Chromatography-Mass Spectrometer (LC-MS) based method to establish retention times of LC-PUFAs and VLC-PUFAs by using standard VLC-PUFAs, 22:6 (DHA), 24:5(n-3), 32:6 (n-3), 34:5 (n-3) and 34:6 (n-3) and mixed VLC-PUFAs extracted from bovine retina as VLC-PUFA standards (see Methods). LC-PUFAs and VLC-PUFAs in samples were extracted using the Bligh-Dyer method and detected using the established retention times in full scan (m/z 250–800) MS under negative mode (**Fig. 1A**, **Table 2**). The identification of each VLC-PUFA (e.g. from bovine retina samples) was achieved through extraction of a chromatographic peak for each of the corresponding [M-H]^-^ ions (**Fig.1B**) in full scan (FS) mode. Then, a separate parallel reaction monitoring (PRM) method was also used to obtain the fragments of different VLC-PUFA peaks for identification. The product ion spectra of the precursor ion [M−H]− of 32:6 and 34:5 at m/z 467.3893 and m/z 497.4368 showed the product ions at m/z 423.4004 and m/z 453.4459 by loss of 44 Da at 35% collision energy (CE), while peak clusters with a mass difference of 14 Da corresponding to CH2 with increased CE. A similar fragmentation pattern was observed from DHA standard (**Fig S1A, B**). The linear range and reproducibility of the FS mode-based quantification were further evaluated by using 32:6 and fatty acid extract, respectively. For 32:6, the linear range was from 50 nmol/L to 1000 nmol/L with a correlation coefficient of 0.994, and the limit of quantification was 5 nmol/L (**Fig. S1C**). The standard deviations of the peak area of the 5 VLC-PUFAs of interest in bovine retina extract were below 15% (n=5) (**Fig. S1D**). Bovine and mouse eyecups, as well as RPE cells isolated directly from eyecups, exhibited a high abundance of VLC-PUFAs ranging from 32:4 to 36:6, and the composition resembled that of retina tissue and POS (**Fig. 1C** and **Fig. S2A**). On the contrary, cultured RPE cells, such as the widely used ARPE-19 cell line, primary bovine RPE cells, and human induced pluripotent stem cell iPSCs-derived RPE have minimal amounts of endogenous VLC-PUFAs (Fig. 1D and Fig. S2B). This suggests that VLC-PUFAs in isolated RPE cells most probably originated from phagocytosed POS *in vivo*. Therefore, the deficiency of VLC-PUFAs in cultured RPE cells suggested the feasibility of using the abundance of VLC-PUFAs after incubation with photoreceptor outer segments as a measure of phagocytosis activity in RPE cells.

### Quantification of phagocytosis in RPE cells

To evaluate the feasibility of the VLC-PUFA-based method for phagocytosis quantification, we followed the workflow presented in **Fig. 2A**. Briefly, ARPE-19 and primary bovine RPE cells were plated at equal density and grown to confluency in 12 well plates. Then, the ARPE-19 and primary bovine RPE cells were exposed to 20 POS/cell for 5 h and 0.5h, respectively, following previously published protocols (8). To terminate phagocytosis, cells were washed 3 times with PBS, trypsinized to remove free and bound POS (22) and collected for analysis. Total fatty acids were extracted and quantified using LC-MS method described above. To assess phagocytic activity, the abundance of VLC-PUFAs 32:6, 34:4, 34:5, 34:6, 36:6 was quantified as markers of internalized POS. All VLC-PUFAs present in POS were detectable in the cells that were challenged with the cargo, *i.e.* in both ARPE-19 cells and primary bovine RPE cells. An apparent increase was observed for PUFAs longer than 30:4, otherwise not detectable in these cells (**Fig. 2B, C**). We therefore concluded that the lipidomic method is able to detect phagocytosed POS in RPE cells.

**Figure 2.**
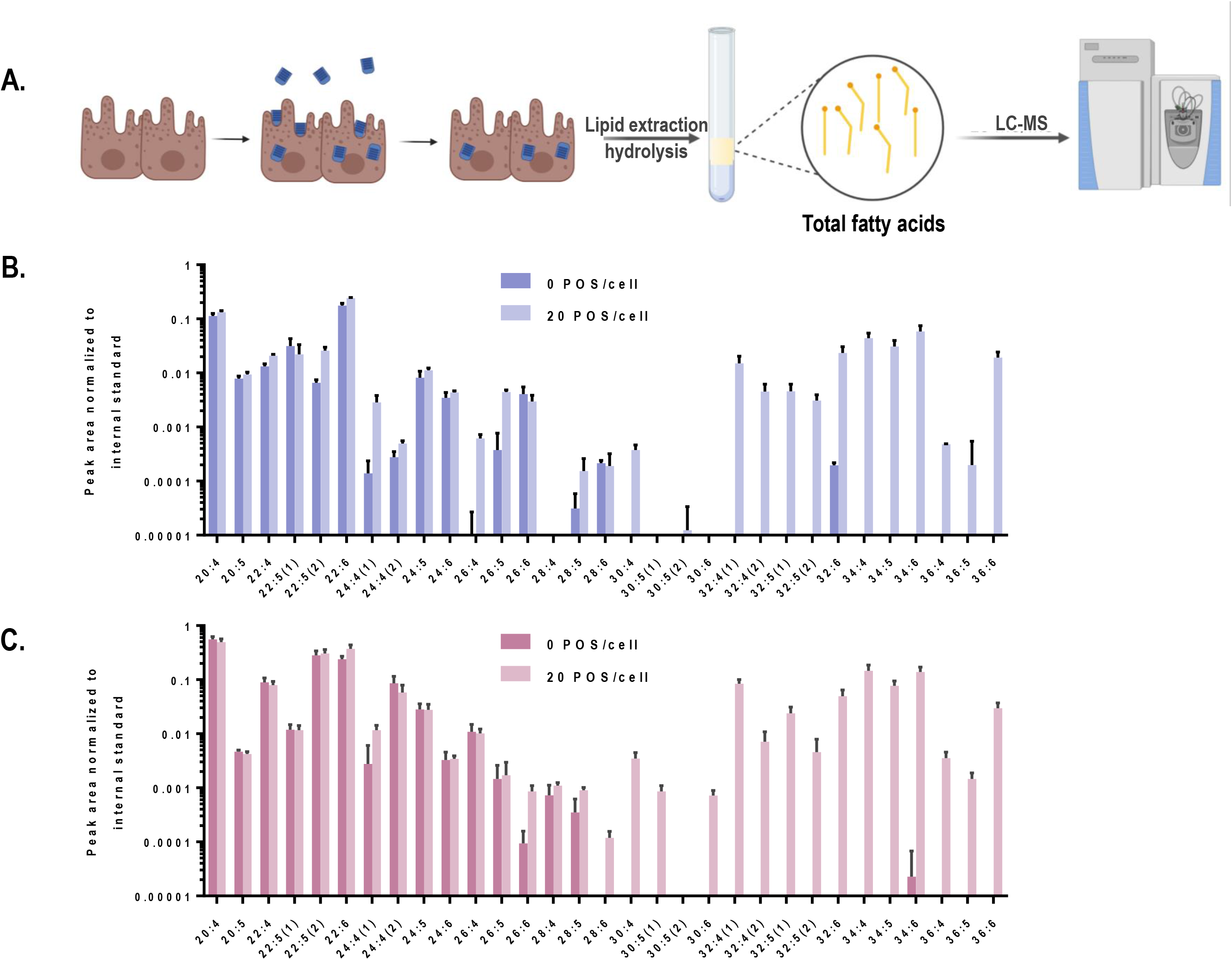
VLC-PUFAs-based quantification of phagocytosed POS in RPE cells. **A.** RPE cells were challenged with unlabelled POS. After washing and trypsinization to remove unbound and surface-bound POS, lipids were purified, fatty acids was extracted and subjected to LC-MS analysis (Created with BioRender.com) LC-PUFA and VLC-PUFA composition of ARPE-19 cells **(B)** and primary bovine RPE cells **(C)** before and after 5 h and 0.5 h incubation, respectively, with POS. Minimal amounts of endogenous VLC-PUFAs were detected in untreated RPE cells, whereas VLC-PUFAs were highly abundant in RPE cells after POS incubation (n=3).

Next, we compared our VLC-PUFA-based assay with the current standard fluorescent phagocytosis assay in ARPE19 cells (19). To do so, ARPE-19 cells were challenged with fluorescently labeled POS (FITC-POS) using an optimized concentration of POS (20 POS/cell) and incubated for 2 and 5 hours. Following standard protocol (8), after fixation and without permeabilization, the cells were briefly stained with an antibody against the C-terminal nine amino acids of bovine rhodopsin (1D4). Internalized FITC-POS appear as green particles on fluorescence microscopy, whereas surface-bound POS appear yellow, due to the overlapping FITC- and secondary antibody (red)-derived fluorescence signals, (**Fig. 3A**). Fluorescence-based quantification showed that the number of engulfed POS increased from 21.01±10.49 to 32.12±8.280 (∼53% increase) from 2 to 5 hours when incubated with 20 POS/cell, (**Fig. 3B, C**). Then we assessed whether the new method is applicable for the detection of the dynamic property of phagocytosis. ARPE-19 cells were challenged with POS with varying POS concentrations (0, 10, 20, 40 POS/cell), and POS uptake by the cells was quantified by the abundance of marker VLC-PUFAs. The levels of 32:6, 34:4, 34:5, 34:6, 36:6, which were not detected in unchallenged ARPE-19 cells, showed gradual, significant increases of VLC-PUFA content after incubation with increasing amounts of POS per cell (**Fig. 3D, and Fig. S3A, B, C**). Similarly, primary bovine RPE cells also showed a significant increase of VLC-PUFAs with increasing amounts of POS per cell and with increasing incubation time (**Fig. 3E, F**).

**Figure 3.**
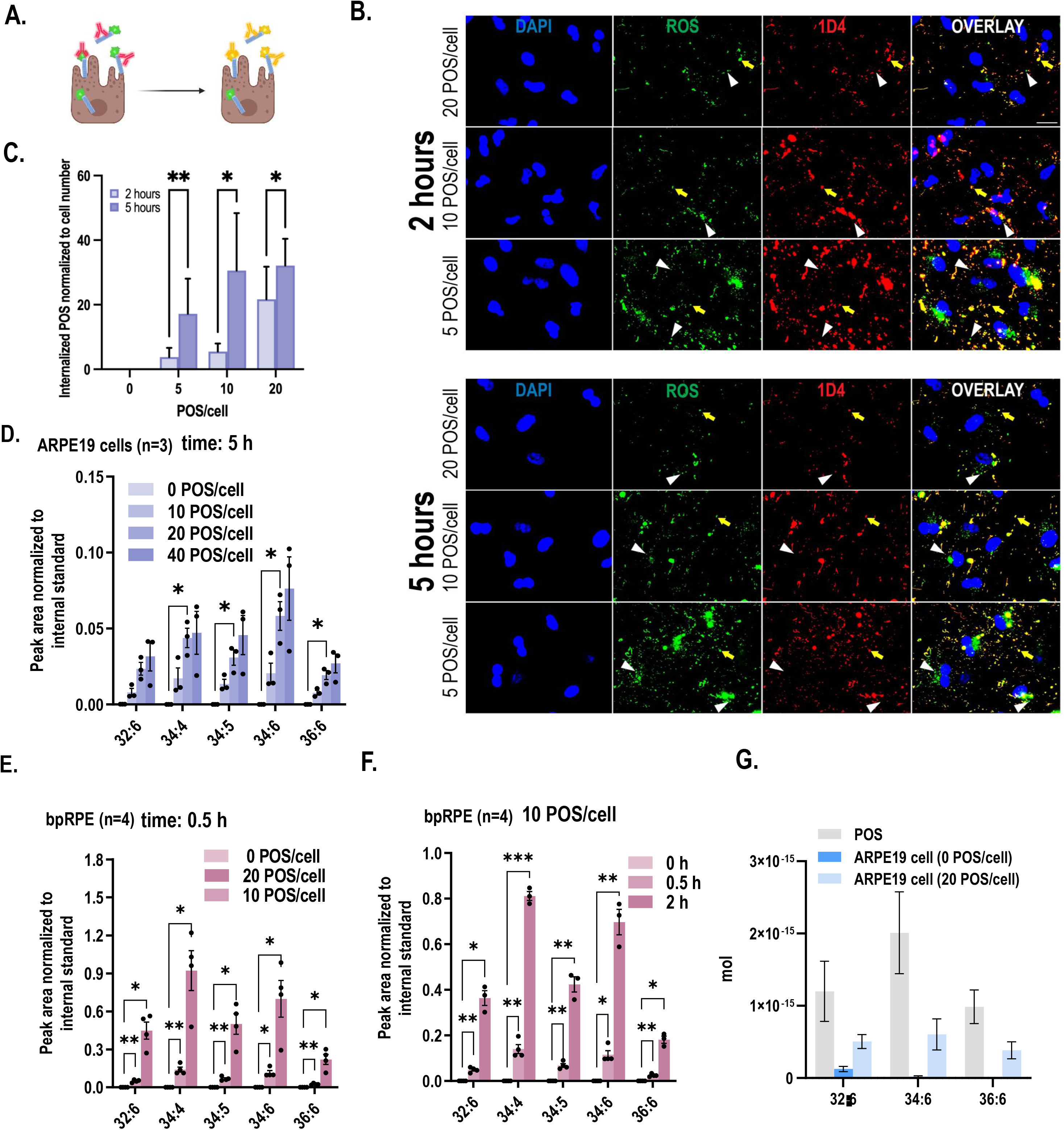
VLC-PUFAs-based quantification can detect gradual changes in phagocytosis. **A.** Fluorescence microscopy can distinguish between surface-bound FITC-POS, which appear yellow in a color overlay of FITC (green) with secondary antibody (red), and internalized POS, which appear green (Created with BioRender.com) **B.** ARPE-19 cells were challenged with FITC-POS for 2 or 5 hours and incubated with antibody against C-terminal epitope of rhodopsin (1D4). Yellow arrows indicate bound POS, while white arrowheads indicate internalized POS, which are only visible in the FITC image. (scale bar = 20μm) **C.** Fluorescence imaging quantification of internalized POS per cell nuclei (n=9 in each group). **D**. Increasing amounts of VLC-PUFAs detected in ARPE-19 cells when challenged with increasing POS concentrations. **E,F.** Increasing amounts of VLC-PUFAs measured in primary bovine RPE cells (bpRPE) when challenged with increasing POS concentrations (**E**) or increasing incubation times (**F**). **G.** Quantification of the amount of 32:6, 34:6, and 36:6 per POS and per cell after 5h of phagocytosis.

Next, using the VLC-PUFA-based approach, we attempted to quantify the amount of the individual VLC-PUFAs per POS and per cell after incubation with POS. The result suggested that there is 1.20E-15 mol, 2.01E-15 mol and 9.86E-16 mol 32:6, 34:6, 36:6 per POS, respectively. After incubation with 20 POS/cell for 5 h, the amount of 32:6, 34:6, and 36:6 in ARPE19 cells increased 3.80E-16 mol, 5.85E-16 mol and 3.83E-16 mol/cell, respectively (**Fig. 3G**). This data shows the level of precision that can be attained when VLC-PUFA based method of quantification is applied.

### Validation of the role of MERTK, GAS6 and MFG-E8 in RPE phagocytosis by VLC-PUFA-based assay

We further evaluated whether the VLC-PUFA-based assay could detect impaired RPE phagocytosis secondary to knockdown of genes known to be key molecules in the binding and engulfment of POS, namely the engulfment receptor MERTK, its ligand GAS6, and the αvβ5 integrin ligand MFG-E8 (23-26). We performed *MERTK/GAS6/MFG-E*8 triple knockdown in ARPE19 cells and detected 50%, 90%, and 60% knockdown of *MERTK*, *GAS6*, and *MFG-E8* mRNAs, respectively (**Fig. 4A**). Next, wildtype and triple knockdown (triple KD) ARPE-19 cells were exposed to 10 POS/cell, and the VLC-PUFA content was measured after 2 hours. The quantification of marker VLC-PUFAs 32:6, 34:4, 34:5, 34:6 and 36:6 decreased 84.82%, 76.63%, 94.70%, 76.50%, 77.40%, respectively, (82.01%±7.90%) in triple KD cells when compared to the control group. The opsin immunoblot-based method indicated a decrease of ∼91% of rhodopsin content (1D4) from internalized POS in triple KD cells (**Fig. 4C, D**). This decrease was consistent with the result obtained by the VLC-PUFA-based assay, indicating the validity of the proposed method.

**Figure 4.**
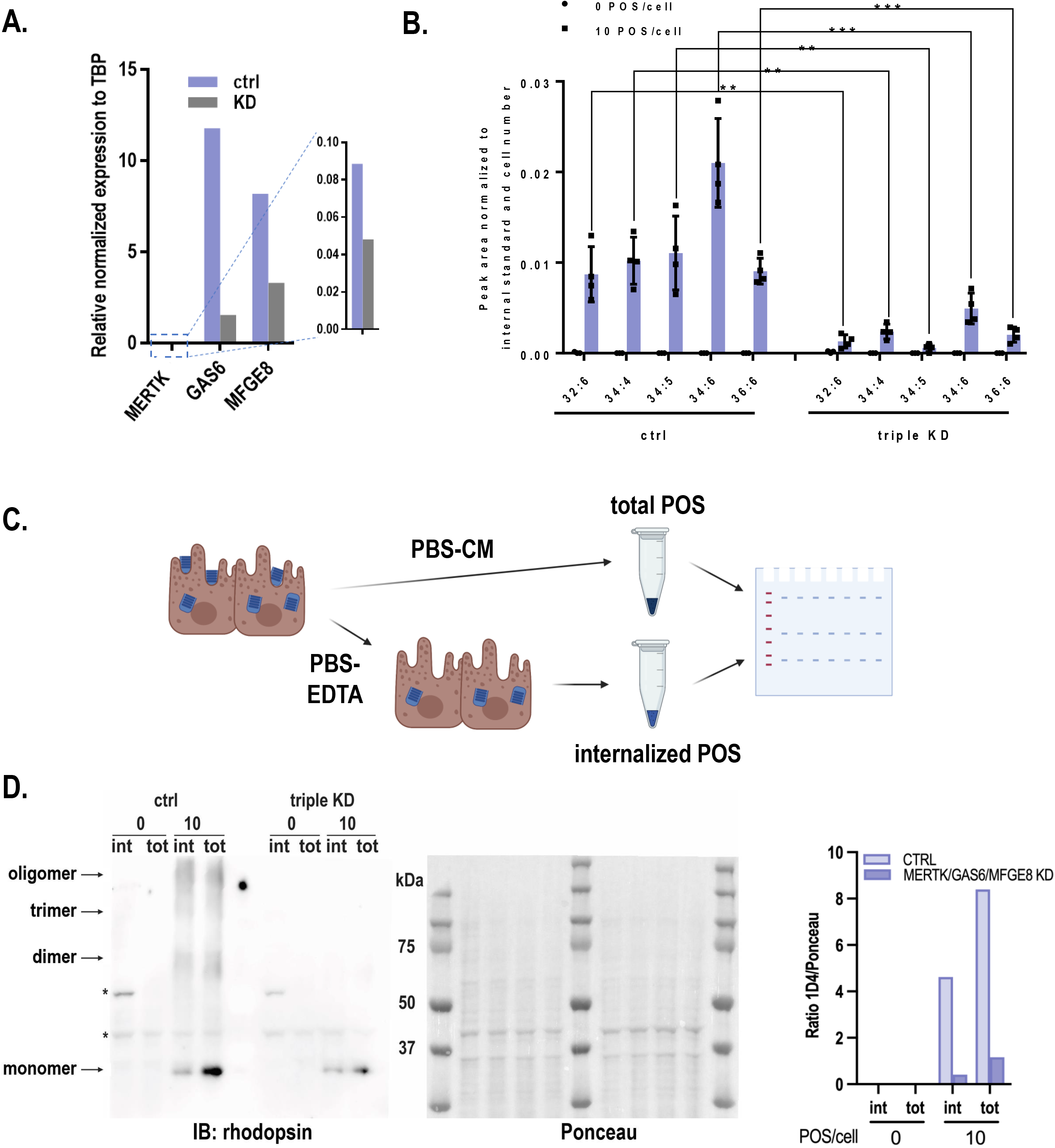
VLC-PUFAs-based quantification can detect impaired phagocytosis. **A.** *MERTK, GAS6, MFGE8* mRNA knockdown efficiency measured by RT-qPCR relative to TBP expression. **B.** Significantly lower levels of VLC-PUFAs detected in triple knockdown ARPE-19 cells challenged with 10 POS/cell for 2 hours. (n=4, ** = p<0.01, *** = p<0.001) **C.** Rhodopsin immunoblotting analysis of POS phagocytosis can distinguish between internalized POS (treated with EDTA before lysis) and total POS (without EDTA treatment). Triple KD cells show less internalized (int) and total (tot) POS after 2 hours **(D**), as quantified by opsin content. Ponceau S staining was used for total protein normalization. * = non-specific bands (n=3)

### Use of VLC-PUFA based phagocytosis assay in a high throughput format

To test the potential usage of the proposed assay as a high throughput screening method, the phagocytosis assay was performed in 48-well plates and assessed for reproducibility of VLC-PUFA detection. In brief, 0.03 x 10^6^ cells were plated, grown to confluency, and incubated with 20 POS/cell for 2 hours. Total fatty acids were extracted, and the reproducibility of 5 independent experiments was analyzed using the method described above. Coefficients of variation for FA 34:4, 34:6 and 36:6 were 16.65%, 16.80% and 19.75%, respectively (**Fig. 5A**).

**Figure 5.**
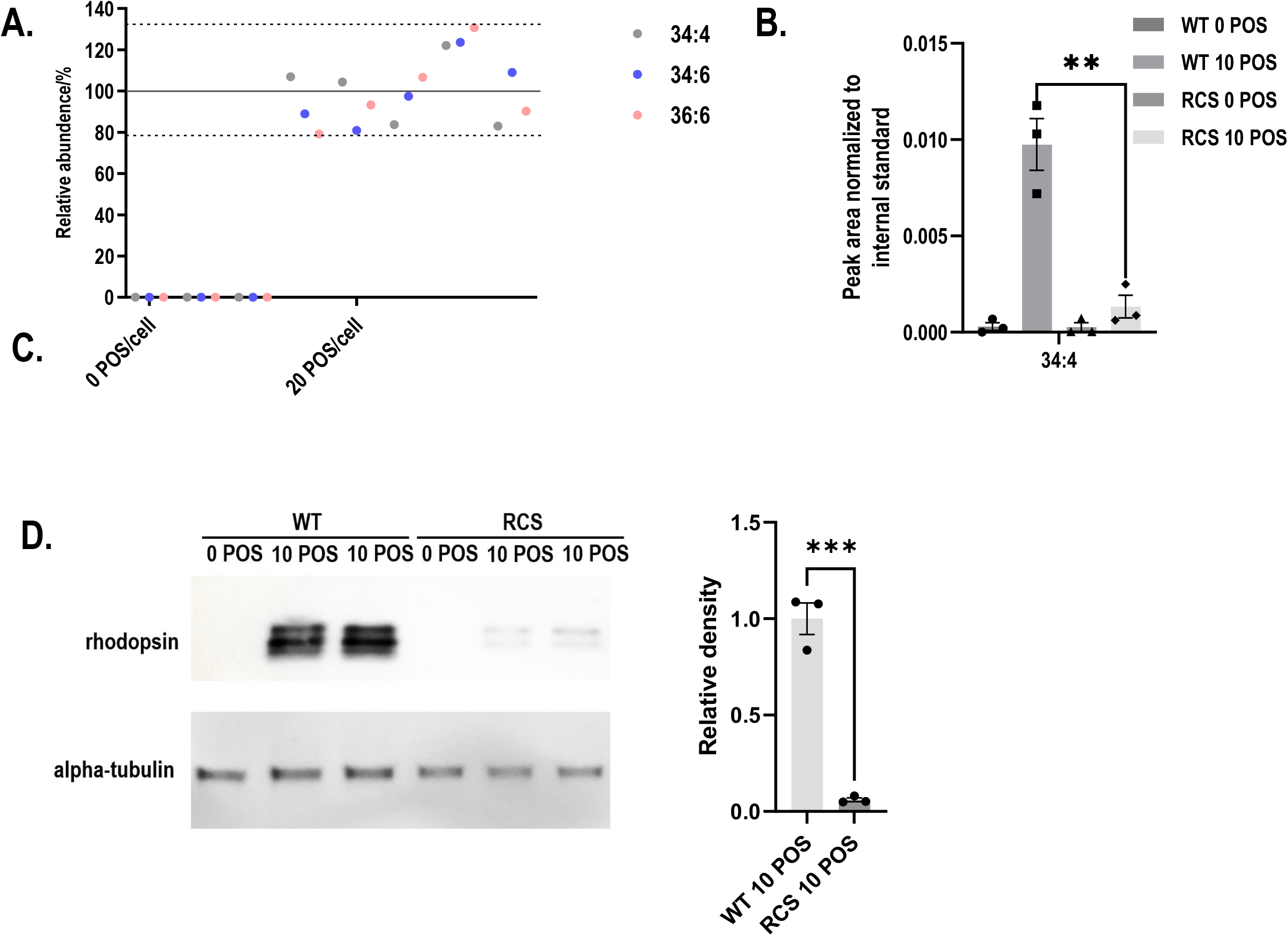
VLC-PUFA based assay allows for the quantification of phagocytosis by RPE cells on 48-well plates. **A.** Reproducibility of the quantification of phagocytosis of ARPE19 cells on 48-well plates showed coefficients of variation <20% for 5 independent experiments. **B.** VLC-PUFA phagocytosis assay on 48-well plates detected impaired phagocytosis of primary RCS cells (n=3). **C.** Rhodopsin immunoblotting analysis of POS phagocytosis detected impaired phagocytosis of primary RCS cells (n=3).

Finally, we used our newly developed assay to compare the phagocytic capacity of primary RPE cells prepared from WT and mutant rats with known phagocytosis competence and deficiency, respectively. The Royal College of Surgeons (RCS) rat displays a defect in RPE phagocytosis as a result of a mutation in the *Mertk* gene(3). It is a common model used to study the development of cellular therapies for retinal diseases. Therefore, we tested our proposed assay on primary RPE cells cultured from WT and RCS rats. Following POS challenge, total fatty acids were extracted using the method described above and quantified using mass-spectrometry. Compared to WT RPE cells, phagocytosis of RCS RPE cells on 48-well plates was reduced 86.4% by 34:4 based quantification (**Fig. 5B**). This result was in line with earlier publications (27) and with western blot analyses performed in parallel (93.9%) (**Fig. 5C**).

## DISCUSSION

Currently, several common techniques are used to study RPE phagocytosis by measuring POS uptake in cultured RPE cells. However, none of them is able to assess the fate of phagocytosed POS lipids. In this paper, we present a novel VLC-PUFA-based approach for quantifying RPE phagocytic function that detects POS uptake by RPE cells at varying POS concentrations and incubation times, offering a robust, quantifiable, and unbiased approach for high-throughput assay for genetic and drug screening.

VLC-PUFAs are abundant in the outer segment of photoreceptors, where they are especially enriched in the center of the photoreceptor discs, in close proximity to rhodopsin (16). This specific localization most probably provides a highly fluid environment for proteins involved in the visual cycle and residing in these membranes. In the process of daily renewal, ∼10% of the rod outer segment is phagocytosed by RPE, therefore enriching the RPE cells in VLC-PUFAs. As mentioned above, VLC-PUFAs are recycled back to the photoreceptors; however, it is not clear whether RPE cells also synthetize VLC-PUFAs endogenously. To investigate this, we first developed an LC-MS based method for VLC-PUFAs quantification. The quantification of VLC-PUFAs remains challenging due to their low abundance and the lack of commercial standards. The procedures most widely used for determining VLC-PUFA levels are Gas Chromatography-Mass Spectrometer (GC-MS) and GC-flame ionization detection following methyl-esterification (9, 15). These methods are highly reproducible and are able to distinguish between n-3 and n-6 PUFAs due to hard ionization capability. However, due to the low volatility of VLC-PUFAs, the method requires the use of special columns and higher temperature for separation on GC columns. On the other hand, soft ionization in Liquid Chromatography-Electrospray ionization-tandem Mass Spectrometer (LC-ESI-MS) is more suitable for identifying larger, more complex compounds since LC-ESI-MS retains precursors during ionization (21) (32). Using this method, we compared the VLC-PUFA profiles of RPE cells isolated directly from the bovine retina, and therefore engaged in the diurnal cycle of phagocytosis, and primary RPE cells, cultured *in vitro* for several days without the presence of outer segments. Our data unequivocally show that, although RPE cells express genes involved in the elongation of long-chain PUFAs (ELOVL2 and ELOVL4), they do not maintain detectable levels of VLC-PUFAs themselves.

Notably, the levels of LC-PUFA and VLC-PUFA detected in bovine retina, RPE and POS are slightly different than that detected in other studies using GC-MS method. It is possible that different methods can have different abilities to ionize and detect different fatty acids. We also believe that our detailed analysis shows us some new information. For example, the levels of 32:6 fatty acid seem to be higher in bovine retina than in bovine POS. In contrast, other VLC-PUFAs have higher abundance in POS, when compared to retina. This data suggests that there might be a separate pool of VLC-PUFA (32:6) which is not located in POS and therefore, the abundance of this FA does not follow the other VLC-PUFAs. The location and role of 32:6 FA in the retina is further investigated in our laboratory using several orthogonal methods.

With this very sensitive method of detection and quantification of VLC-PUFAs in hand, we then applied this method as a tool for quantifying phagocytosis activity. Indeed, using isolated bovine POS and several different incubation conditions, we could detect significant increases in VLC-PUFA content in RPE cells after POS challenge, which we could confirm with established standard assays. Therefore, our new approach can be used as an orthogonal method of quantification that is complementary to the observations obtained by other assays.

Interestingly, the VLC-PUFA-based method seems to be more sensitive than the fluorescence-based assay when quantifying POS uptake by ARPE-19 cells incubated with 20 POS/cell. Although the two methods both measured an increased phagocytic trend when increasing the POS number, the new VLC-PUFA-based quantification allows for absolute quantification of the VLC-PUFAs in single POS, as well as single cell after incubation with POS. It is possible that, considering the high sensitivity and wide linear range of Mass Spectrometry, this VLC-PUFAs-based quantification may represent the optimal method to measure a large range of phenotypes and defects in phagocytosis. For example, due to the different processing rates for lipids vs other components of POS *e.g.,* proteins, the new method provides an unprecedented opportunity to study phagocytosis in a more measurable way on the level of individual lipid in the future.

Finally, we decided to test whether the VLC-PUFA-based assay is able to detect defects in phagocytosis caused by lack of key regulators of the process. Mer tyrosine kinase (MERTK) is essential for the efficient engulfment of POS by RPE (23) and mutations in *Mertk* have been identified in patients with *retinitis pigmentosa* (3, 9, 28). Growth arrest-specific protein 6 (GAS6) specifically and selectively stimulates phagocytosis by cultured rat RPE cells by ligating RPE surface MERTK, leading to receptor phosphorylation and activation (24, 25). Milk fat globule-EGF 8 (MFG-E8), which localizes to the subretinal space in the retina, acts as bridge ligand in the retina that connects externalized phosphatidylserine of outer segment tips with the αvβ5 integrin POS recognition receptors of the RPE(29). Extracellular MFG-E8 stimulates POS phagocytosis by the RPE *in vivo* and by RPE cells in culture (26, 30). MFG-E8-deficient cultured primary RPE showed impaired binding and, secondarily, engulfment of isolated POS(22). Triple knockdown of *MERTK*, *GAS6*, and *MFG-E8* in ARPE-19 cells impaired phagocytosis, likely by causing defects in both αvβ5 integrin-dependent recognition/binding and MERTK-dependent internalization steps. By using the proposed VLC-PUFA-based assay, we showed a robust decrease of VLC-PUFAs in triple KD cells when compared to the control group. This new method is directly quantifiable, unbiased, and is especially sensitive, due to the application of mass spectrometry. Also, since different VLC-PUFAs are used for quantification in one sample, quantification results can be presented for each VLC-PUFA, or only one FA can be used for a larger number of samples. Moreover, the proposed assay showed reproducible and sensitive detection of VLC-PUFAs, which allowed for the quantification of phagocytosis of RPE cells on 48-well plates. Therefore, the proposed phagocytosis assay enables high-throughput studies of POS uptake or POS lipid processing, including pharmacological or genetic screening, combined with the application of Ultra Performance Liquid Chromatography (UPLC) and high-resolution Mass Spectrometry.

In sum, we present a highly adaptable method for unbiased quantification of RPE phagocytosis through LC-MS and detection of VLC-PUFAs present in POS. The next challenge is to apply this quantitative method for assessment of POS lipids *in situ*.

## ACKNOWLEDGMENTS

Authors would like to thank Dr. Huajun Yan for providing primary bovine RPE cells for experiments.

## AUTHOR CONTRIBUTIONS

DS-K—Conceptualization, data analysis, writing, editing, providing funding; SCF—Conceptualization, editing, providing funding; SAL—experiments, data analysis; FG—experiments, data analysis, writing, editing; ET—experiments, data analysis, writing, editing: FG and ET—equally contributed.

---

**Supplementary Figure 1.**
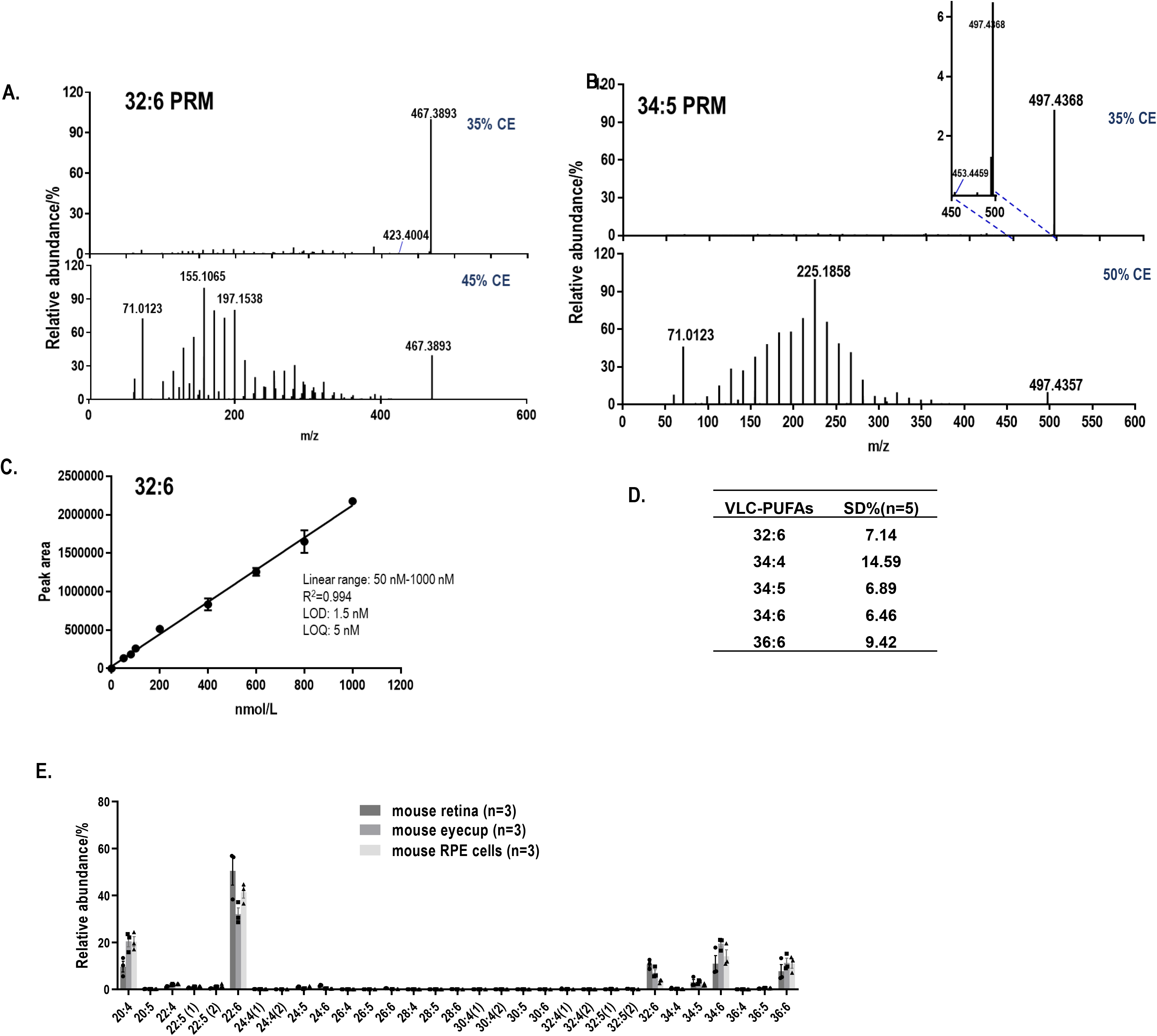
Identification and quantification of VLC-PUFAs by LC-MS and LC-MS/MS method. Peak clusters with a mass difference of 14 Da corresponding to CH_2_ in parallel reaction monitoring (PRM) fragment ion chromatograms of 32: 6 **(A)** and 34:4 **(B)** from bovine retina extract. **C.** Linearity and linear range for 32:6 in FS mode. Linear correlation coefficient = 0.994 from 50 nmol/L to 1000 nmol/L, and the limit of quantification of 32:6 = 5 nmol/L (n=5). **D.** The standard deviations of the peak area of the given VLC-PUFAs from bovine retina extract were below 15% (n=5). **E.** Abundance plots present the relative abundances within the total abundance of all the detected LC-PUFAs and VLC-PUFAs in mouse samples. LC-PUFAs and VLC-PUFAs composition in mouse retina, eyecup and RPE cells exhibited high abundance of VLC-PUFAs, ranging from 32:6 to 36:6. n represents number of retinas and eyecups was used, while in mouse RPE cells sample, n represented number of mice (2 eyecups/mice). Retinas and eyecups were collected in the morning.

**Supplementary Figure 2.**
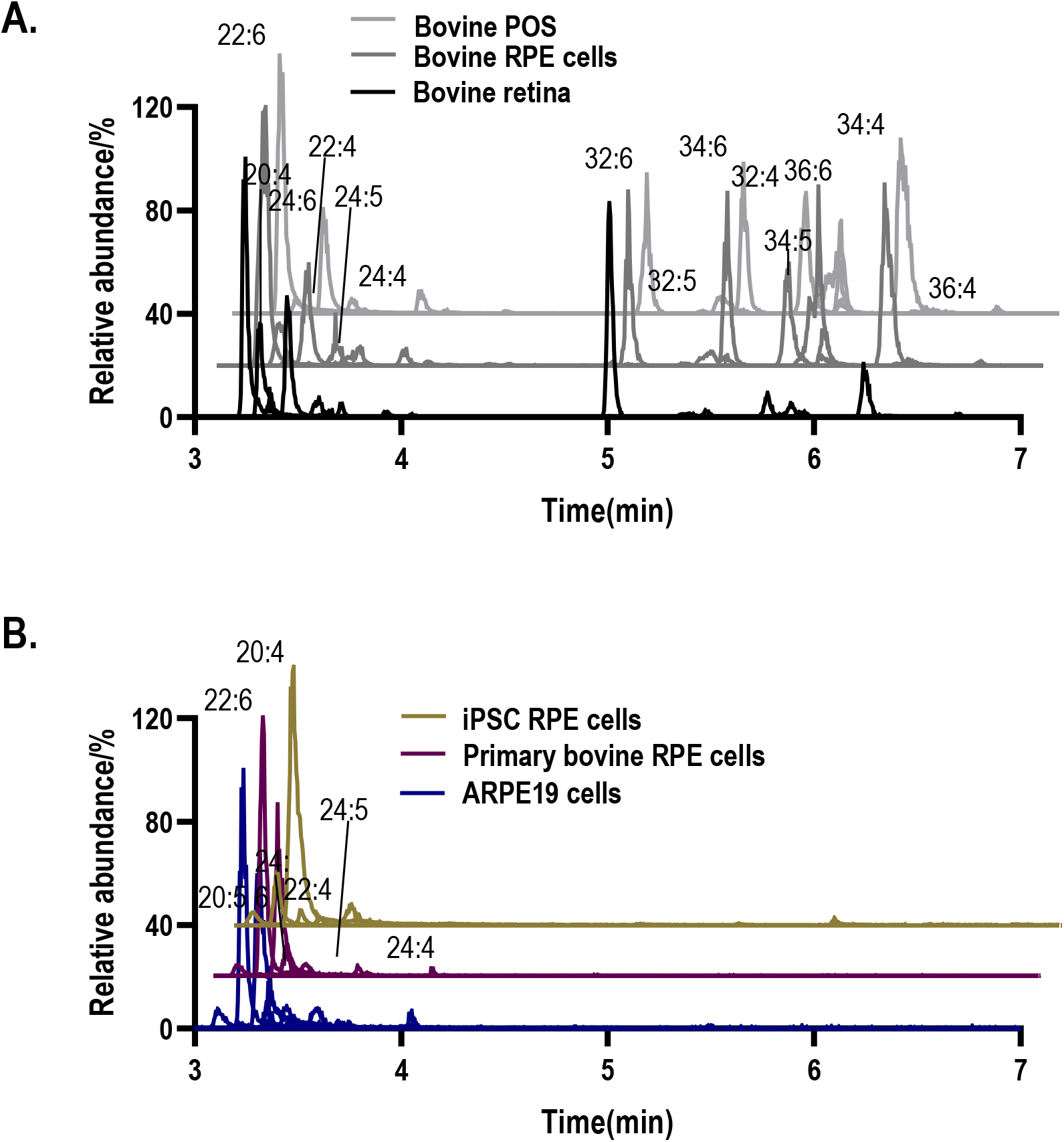
LC-MS chromatograms of LC-PUFAs and VLC-PUFAs. A. EIC of LC-PUFAs and VLC-PUFAs extracted from bovine retina, bovine RPE cells and bovine POS. B. EIC of LC-PUFAs and VLC-PUFAs extracted from ARPE19 cells, primary bovine RPE cells and iPSC-RPE cells.

**Supplementary Figure 3.**
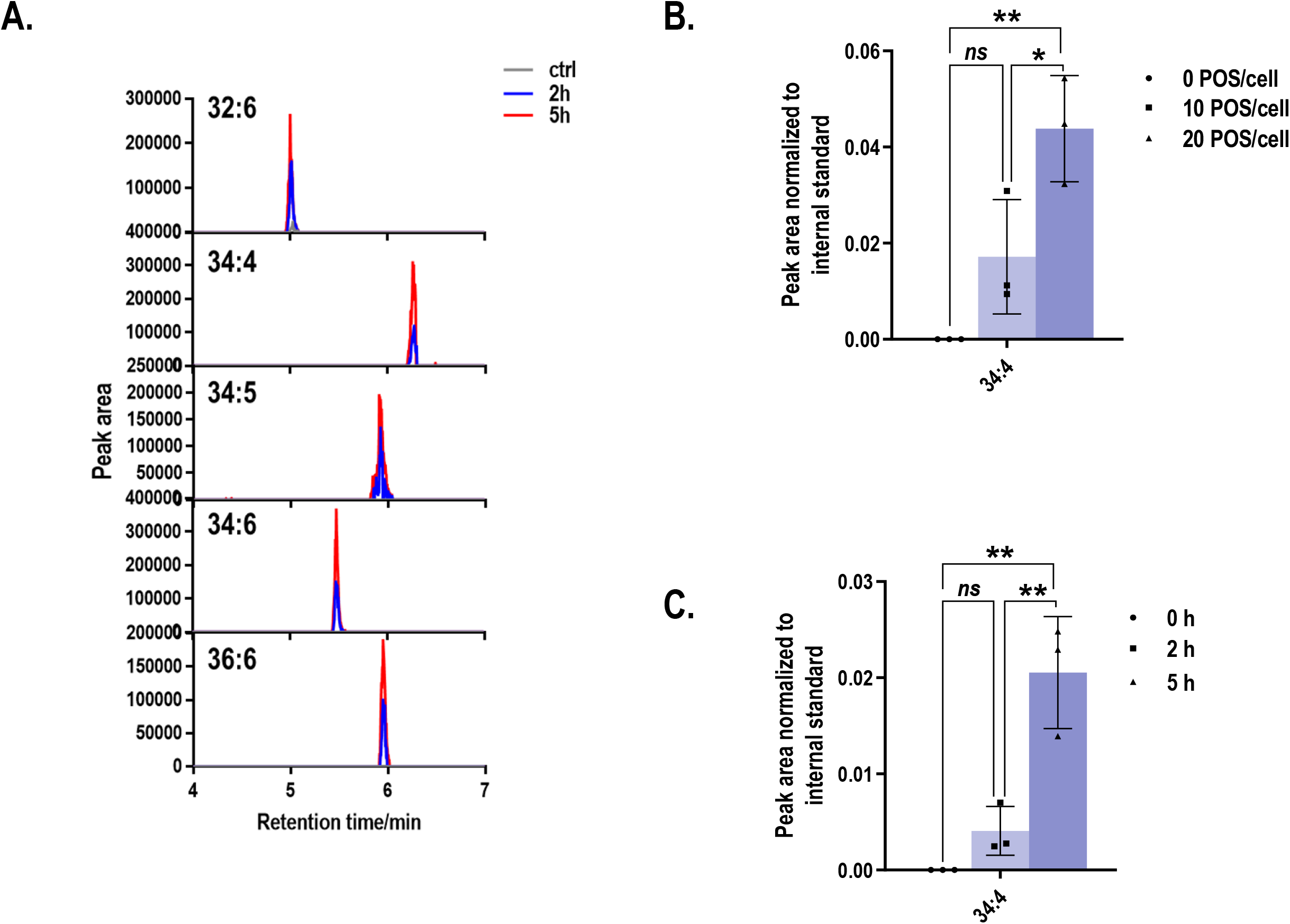
VLC-PUFAs-based quantification can detect gradual changes in phagocytosis. **A.** Extracted ion (EIC) mass chromatograms of marker VLC-PUFAs obtained in FS mode by LC–ESI(-)– MS (FS) after incubating with 20 POS/cell for 2 and 5 hours. Quantification of phagocytosis was performed by using 34:4 as one of POS marker VLC-PUFAs when ARPE-19 cells were challenged with increasing POS concentrations **(B)** or increasing incubation times **C.** (n=3, * = p<0.05, ** = p<0.01)

